# Effect of translation enhancing nascent SKIK peptide on the arrest peptides containing consecutive Proline

**DOI:** 10.1101/2024.02.28.582505

**Authors:** Yuma Nishikawa, Riko Fujikawa, Hideo Nakano, Takashi Kanamori, Teruyo Ojima-Kato

## Abstract

Ribosome arrest peptides (RAPs) such as SecM arrest peptide (SecM AP) and WPPP with consecutive Pro residues, are known to induce translational stalling in *Escherichia coli*. We demonstrate that the translation enhancing SKIK peptide tag, which consisted of four amino acid residues Ser-Lys-Ile-Lys, effectively alleviate translational arrest caused by WPPP. Moreover, the proximity between SKIK and WPPP significantly influences the extent of this alleviation, observed in both PURE cell-free protein synthesis and in vivo protein production systems, resulting in a substantial increase in the yield of proteins containing such RAPs. Furthermore, we unveil that nascent SKIK peptide tag and translation elongation factor P (EF-P) which alleviate ribosome stalling in consecutive-Pro rich protein, synergistically promote translation. A kinetic analysis based on the generation of super folder green fluorescent protein under in vitro translation reaction reveals that the ribosome turnover is enhanced by more than 10-fold when the SKIK peptide tag is positioned immediately upstream of the SecM AP sequence. Our findings provide valuable insights into optimizing protein production processes, which are essential for advancing synthetic biology applications.

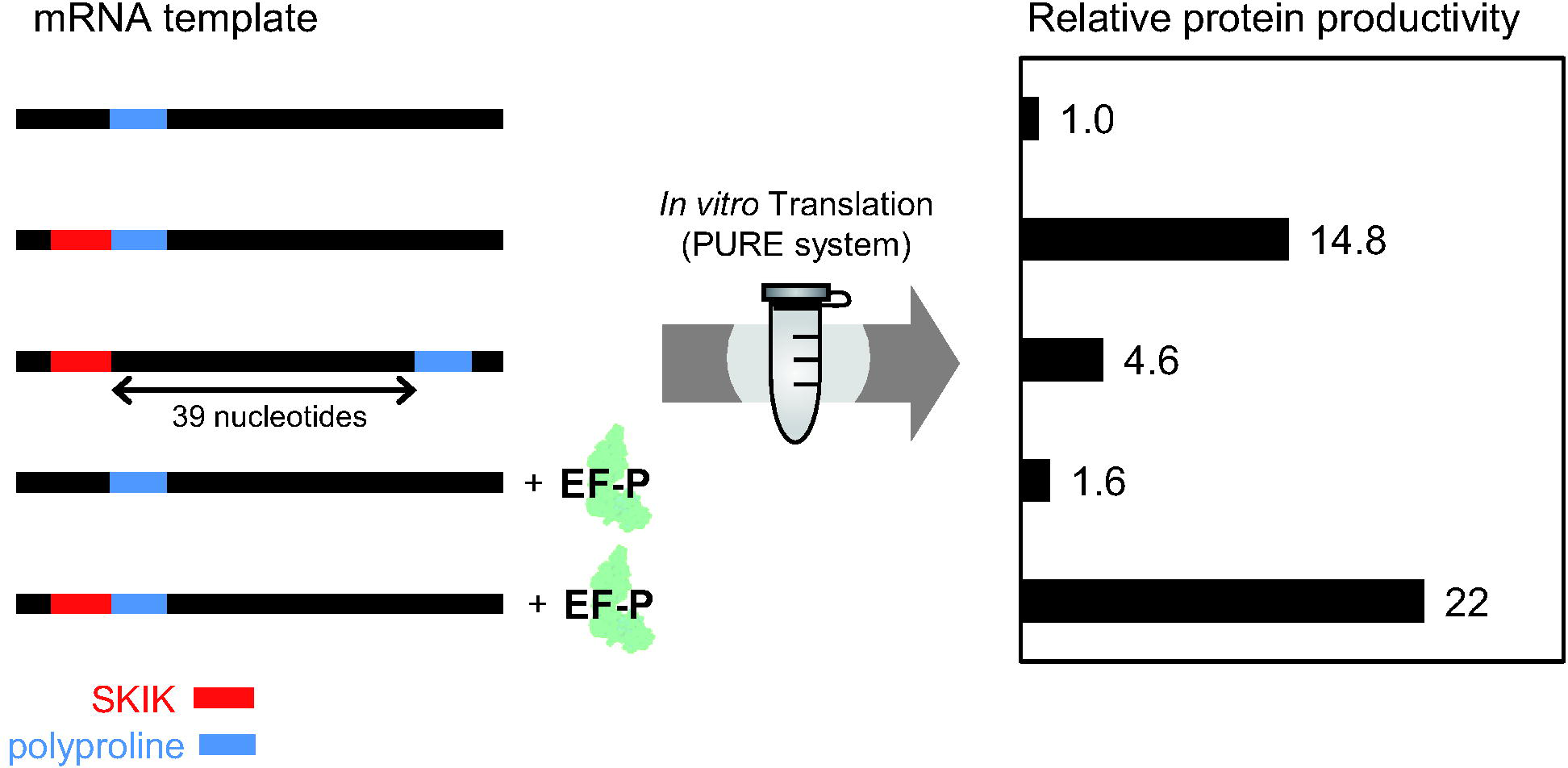

## Introduction

Protein synthesis is a fundamental process in synthetic biology, and the translation by ribosomes is an essential step in protein synthesis. Nonetheless, achieving high productivity for all proteins of interest remains a challenge, often leading to encounters with difficult-to-express proteins for reasons yet to be fully elucidated. Addressing this issue necessitates considering various strategies. These include codon optimization^1–6^, loosening the secondary structure of mRNA^7–9^, utilizing a combination of host-vector systems comprising ribosome binding site (RBS), promoter, and terminator^10–13^. Furthermore, co-expression of molecular chaperons^14^, and the use of fusion protein tags (e. g., maltose binding protein domain, glutathione S-transferase, small ubiquitin-related modifier) have proven beneficial^15,16^. optimizing culture conditions plays a pivotal role in enhancing productivityl^17–19^.

Recently, some nascent polypeptides generated during the translation process have been reported as ribosome arrest peptides (RAPs), which cause translational stalling by interacting with internal components of the ribosome when they pass through the ribosome exit tunnel in the nascent state. RAP-mediated translation arrest is directed by the amino acid sequence of the nascent polypeptide chain instead of the mRNA nucleotide sequence. It is independent of codon usages or the secondary structure of mRNA^21^.

RAPs have been found in a variety of organisms, including SecM and CmlA leader in *E. coli*^20–23^, VemP in *Vibrio alginolyticus*^24,25^, and MifM in *Bacillus subtilis*^26^. Among them, SecM is the most studied, and the partial sequence F^150^XXXXWIXXXXGIRAGP^166^ (X is any amino acid) at the C-terminus is required and sufficient for causing ribosomal stalling^27^, where new peptidyl bond formation is inhibited when the 166th Pro is going to be synthesized (P-site: SecM_1-165_-Gly-tRNA, A-site: Pro-tRNA) at the peptidyl-transferase center (PTC). Due to the structural rigidity and unique status as an N-alkylamino acid, Pro-tRNA at the A site slows down the peptidyl bond formation in the RAPs containing SecM and poly-Pro^28^. As such, the WPPP (FXXYXIWPPP: X is any amino acid) sequence, artificially created RAP, efficiently induces ribosomal stalling due to its consecutive Pro residues^29,30^. The universally conserved bacterial elongation factor called elongation factor P (EF-P, PDB 1UEB) is responsible for preventing the translational stalling caused by consecutive Pro in prokaryotic cells^31–34^. The shape and size of EF-P are similar to tRNA and it interacts with the ribosome via the E site in the PTC^35–37^. However, Woolstenhulme et al. showed that EF-P had little or no effect on stalling by the full-length WPPP AP (FXXYXIWPPP) and RXPP motif^30^. We have previously reported that the insertion of an “SKIK peptide tag” composed of the four amino acids Ser-Lys-Ile-Lys at the N-terminus of difficult-to-express proteins, enhances protein production not only in both *E. coli* in vivo and in vitro systems but also in yeast^38^. More recently, we revealed that changing nucleotide sequences corresponding to these four amino acids had no effect on protein production and that the nascent SKIK peptide generated during the translation process could enhance the translation process, but not transcription^39^. Surprisingly, ribosomal stalling by RAPs such as SecM AP (FSTPVWISQAQGIRAGP) and chloramphenicol induced CmlA leader (KNAD) was canceled by a nascent SKIK peptide generated immediately upstream of the RAPs. Moreover, the combined investigation of the SKIK peptide tag and SecM AP suggested that the SKIK peptide, even when positioned internally within the protein, could enhance translation. Additionally, it was suggested that the distance between SKIK and the AP motif might be crucial for SKIK to effectively alleviate translational stalling induced by RAPs. However, it should be noted that in our previous study, the SKIK peptide tag showed no significant impact on translation when it was positioned immediately upstream of the WPPP AP (FQKYGIWPPP)^39^. Based on our recent findings, we hypothesized that ribosomal stalling induced by RAPs containing a poly-Pro motif could be alleviated by optimizing the distance between the nascent SKIK peptide and the poly-Pro motif. In this study, we investigated the positional effects of the SKIK peptide tag on the ribosomal stalling caused by the poly-Pro AP motif in both *E. coli* in vitro and in vivo systems. Additionally, we uncovered that EF-P and the SKIK peptide tag synergistically enhance translation, resulting in a significant increase in protein production. Translation promotion was observed only when the SKIK peptide was tagged, not by adding the synthesized free SKIK peptide to the in vitro translation system, emphasizing the importance of the nascent state of the SKIK peptide in counteracting arrest. Furthermore, we performed a kinetic analysis using an in vitro translation system to elucidate how the SKIK peptide tag influences on translation.

## Results and Discussion

### Influences of the distance between SKIK and WPPP arrest peptide

As described above, the ribosomal stalling by WPPP (FQKYGIWPPP), an artificial AP containing consecutive Pro, was not affected by the SKIK peptide tagging while the ribosomal stalling by SecM AP was more effectively reduced when the SKIK peptide tag was positioned closer^39^. We therefore created DNA constructs corresponding to the amino acid sequence depicted in Figure 1 to explore the hypothesis that an optimal position for the SKIK peptide tag exists concerning the poly-Pro motif. Here, the SKIK tag positioned immediately before the reported AP motif (FQKYGIWPPP) of WPPP was designated as the standard (+0 aa). Minus and plus numbers indicate the count of deleted amino acid residues from F in the WPPP AP motif and inserted G, respectively. To confirm that only the four amino acids “WPPP” function as an arrestor, a version with -6 aa (SKIKWPPP), where the SKIK moiety was substituted with GGGG or AAAA, was also constructed. In the results of in vitro translation using mRNA, the fluorescence intensity, reflecting the synthesized sfGFP quantity after 90 minutes of reaction, demonstrated a rising trend as the distance between SKIK and WPPP decreased (Figure 2A). Particularly noteworthy was the fluorescence intensity at -5 aa (28,860), where SKIK and WPPP AP were brought closer, being approximately 5 times higher than at +0 aa (5,660), indicating the effective alleviation of ribosomal stalling induced by poly-Pro by the SKIK peptide tag. Western blotting analysis for His tag detection also revealed an increase in the full-length protein synthesis as the distance between SKIK and poly-Pro decreased. Moreover, insertions of G at +4–7 aa and +13 aa also exhibited approximately twice the fluorescence intensity of +0 aa, suggesting the possibility of multiple conditions where SKIK can effectively mitigate the ribosomal stalling caused by poly-Pro. It is worth noting that the fluorescence intensity for -6 aa (SKIKWPPP) was 20,200, whereas GGGG (sequence GGGGWPPP) and AAAA (AAAWPPP) exhibited significantly lower intensities. This indicates that the four amino acids WPPP alone can induce translational stalling, and SKIK counteracts this stalling. In the RAP employed in this study, the WPPP AP motif (FQKYGIWPPP), it was reported that the presence of F, Y, I, and W upstream of PPP is essential for translational arrest. However, Elgama et al. demonstrated that the strength of translation arrest induced by a poly-Pro motif is influenced by the specific amino acid residues immediately upstream^40^. This may support our findings that SKIK has the potential to alleviate translational arrest induced by poly-Pro, and its effectiveness seems to depend on the distance between SKIK and poly-Pro motif.

**Figure 1.**
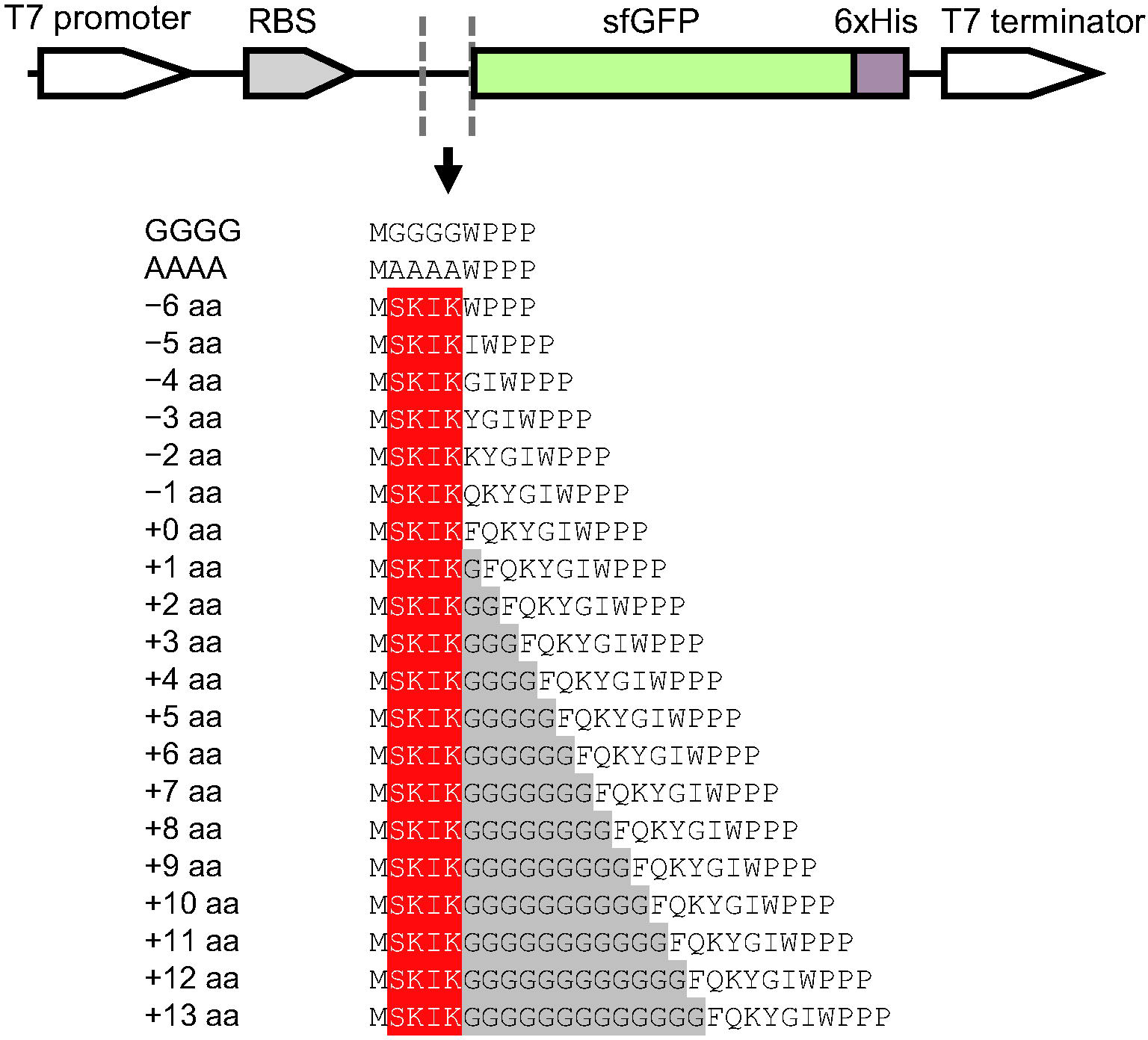
DNA constructs used to study the distance between the SKIK peptide tag and poly-Pro motif of WPPP. The amino acid sequences including start M encoded immediately upstream of sfGFP are shown. SKIK and inserted G are shown in red and gray, respectively.

**Figure 2.**
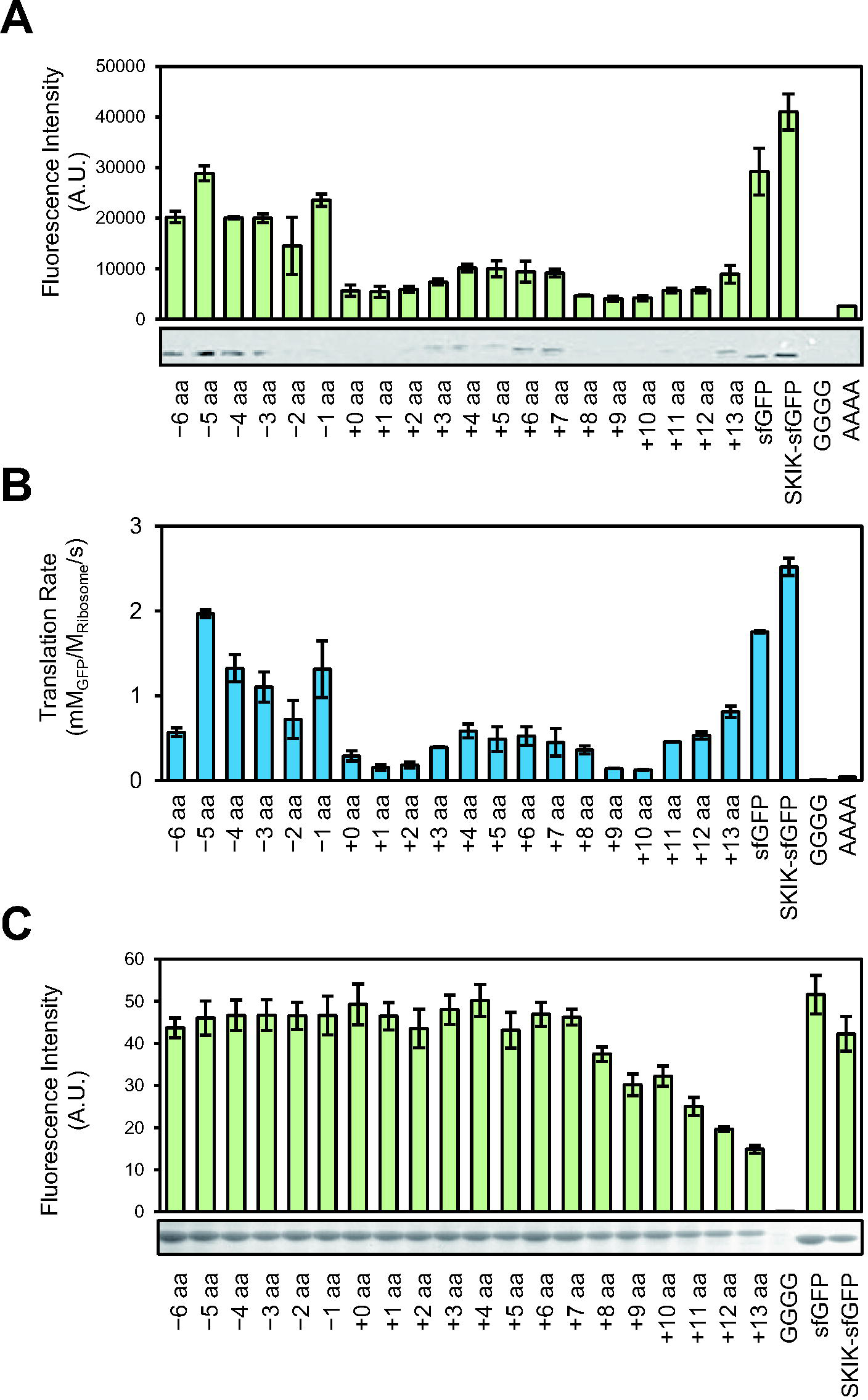
Influences of the distance between SKIK peptide and WPPP poly-Pro motif on protein production and translation rate. The sample names and sequences are corresponding to those of Figure 1. A) Fluorescence intensity and Western blotting analysis of sfGFP detecting His tag. In vitro translation was carried out using mRNA templates (1.66 µM) for 90 min (n = 2). B) Translation rate calculated from the real-time monitoring of the fluorescence intensity of the generated sfGFP during in vitro translation (n = 2, mRNA concentration; 1.66 µM). C) Fluorescence intensity of the produced sfGFP in *E. coli* BL21(DE3) living cell expression system (n = 6). SDS-PAGE and CBB staining analysis of the sfGFP bands are shown.

The translation rate, determined by monitoring the real-time synthesis of sfGFP from mRNA, correlated with the amount of the protein obtained after 90 minutes of reaction (Figure 2A, B). Hence, under the 90-minute CFPS conditions in this experiment, it can be inferred that the final protein amount is indeed influenced by the translation rate. Nevertheless, as the distance increased beyond “+8 aa,” a declining trend in synthesis levels was observed. Upon comparison with the protein production levels of “GGGG,” a sequence excluding SKIK, it can be inferred that translational arrest was alleviated in all sequences containing SKIK, albeit with varying degrees.

The more prominent trend of translational arrest alleviation observed in vivo may be attributed to the fact that in vitro reactions were only 90 minutes, while in vivo, we conducted longer expression induction. Additionally, the presence of natural EF-P in the living cells could contribute to this difference.

To the best of our knowledge, there have been no reports on function of the SKIK peptide to alleviate the ribosomal stalling by poly-Pro or on the importance of their positional relationship. Therefore, our findings will provide a critical and novel insight in the field of synthetic biology.

### Kinetic analysis of the protein synthesis

We then performed a kinetic analysis of translation to better understand the effect of the SKIK peptide tag on translation. Here, SecM AP-sfGFP with and without the SKIK peptide tag and simple translation reaction model were used (Figure 3A, B). For this analysis, we designate mRNA as “Substrate (S),” sfGFP as “Product (P),” and ribosome as “Enzyme (E)” for convenience. We applied the following three assumptions to simplify the analysis: 1) there is no folding rate limitation for sfGFP, 2) there are sufficient elements necessary for translation, and no molecules inhibit the translation reaction, and 3) mRNA and ribosome do not dissociate until the translation of single protein molecule is completed.

**Figure 3.**
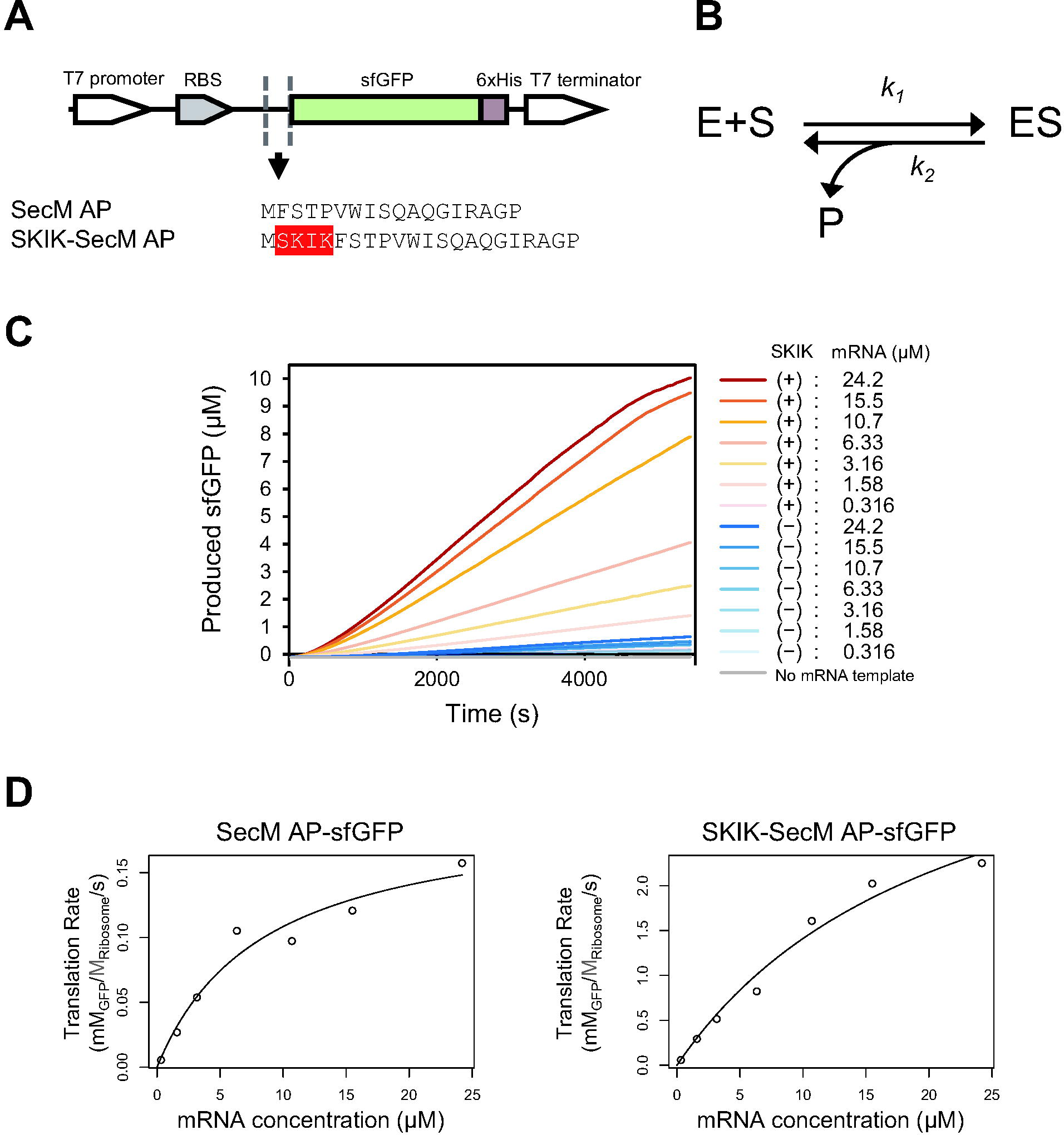
Kinetic analysis of translation by PURE cell-free protein synthesis. A) mRNA constructs and amino acid sequences of SecM AP and SKIK-SecM AP. B) Translation reaction model: E, S, and P represent the ribosome, mRNA, and sfGFP (product), respectively. *k*_1_ and *k*_2_ denote the rate constants for the reactions indicated by the respective arrows. The ES complex encompasses all stages from initiation to termination of translation. C) Real-time fluorescence intensity measurements. Generation of sfGFP of various mRNA concentrations was monitored during 90min (5,400 s) CFPS reaction. Plus and minus in the parentheses indicate the presence and absence of the SKIK peptide tag, respectively. D) Non-linear regression analysis of translation rate.

The translation rate *v* was defined as follows.

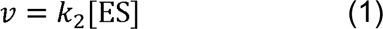

The following equation is given by the steady-state approximation.

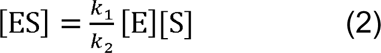

On the other hand, [E] can be expressed as follows:

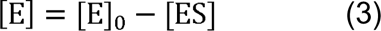

The following equation can be derived by substituting (3) into (2).

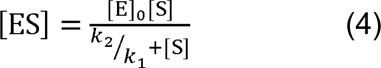

Based on the assumption that k_2_[£]_O_ = V_max_, substituting equation (1) into (4) results in a Michaelis-Menten-like equation as follows:

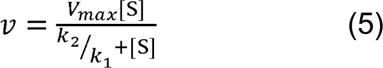

Where [S] = ^k^_2_/_k1_ when the translation rate is 1/2 *V*_max_.

The translation rate was assessed by continuously monitoring the fluorescence intensity of sfGFP generated by CFPS using mRNA templates ranging from 0.316 to 24.2 µM. As shown in Figure 3C, the rate of sfGFP production escalated proportionally with the quantity of mRNA template utilized. Based on these measurements, the translation rate was calculated (Table 1). It was observed that the translation rate increased by over 10-fold when the SKIK peptide was tagged, regardless of mRNA concentrations, peaking at a maximum enhancement of 16.8-fold at 15.5 µM.

**Table 1.** Translation rate.

| mRNA ($\mu\text{M}$) | $v$ ($\text{mM}_{\text{GFP}}/\text{M}_{\text{Ribosome}}/\text{s}$) | | $V_{\text{SKIK}}/V_{\text{No tag}}$ |
| --- | --- | --- | --- |
|  | No tag | SKIK |  |
| 24.2 | 0.157 | 2.25 | 14.3 |
| 15.5 | 0.121 | 2.02 | 16.8 |
| 10.7 | $9.73 \times 10^{-2}$ | 1.61 | 16.5 |
| 6.33 | 0.105 | 0.823 | 7.82 |
| 3.16 | $5.38 \times 10^{-2}$ | 0.514 | 9.56 |
| 1.58 | $2.69 \times 10^{-2}$ | 0.292 | 10.9 |
| 0.316 | $5.48 \times 10^{-3}$ | $5.72 \times 10^{-2}$ | 10.4 |
| No template | $9.52 \times 10^{-4}$ | | |

Utilizing the translation rates calculated for each mRNA concentration in Table 2, nonlinear regression analysis was conducted using the R package “renz”^41^(Figure 3D). The regression curve aptly fits the experimental data, enabling the calculation of, *V_max_* and *k_2_*/*k_1_* based on Equation (5). As a result, the *V_max_* values for SKIK peptide tag absence and presence were determined to be 0.20 mM_GFP_/M_Ribosome_/s and 4.50 mM_GFP_/M_Ribosome_/s, respectively. The addition of SKIK peptide tag led to a 22.5-fold increase in *V_max_*.

**Table 2.** Parameters on translation calculated based on real-time translation monitoring.

| | $V_{\max}$<br>($\text{mM}_{\text{GFP}} \cdot \text{M}_{\text{Ribosome}}^{-1} \cdot \text{s}^{-1}$) | $k_2/k_1$<br>($\mu\text{M}$) | $k_1$ | $k_2$ |
| --- | --- | --- | --- | --- |
| No tag | 0.20 | 8.45 | $2.37 \times 10^7$ | $0.20 \times 10^3$ |
| SKIK | 4.50 | 21.9 | $20.5 \times 10^7$ | $4.50 \times 10^3$ |
| SKIK/No tag<br>(-fold) | 22.5 | 2.59 | 8.65 | 22.5 |

$\text{M}_{\text{GFP}} \cdot \text{M}_{\text{Ribosome}}^{-2} \cdot \text{s}^{-1} \cdot \text{M}_{\text{mRNA}}^{-1}$ and $\text{M}_{\text{GFP}} \cdot \text{M}_{\text{Ribosome}}^{-2} \cdot \text{s}^{-1}$ , respectively. $\text{M}_{\text{Ribosome}}$ and $\text{M}_{\text{mRNA}}$
represent the concentrations of ribosome and mRNA in mol/L, respectively.

The calculated *k_2_*/*k_1_* values were 8.45 µM without SKIK and 21.9 µM with SKIK, representing a 2.59-fold increase upon SKIK peptide tag addition (Table 3). [E]₀ = 1 µM and *V_max_* = *k_2_* [E]₀ were used to calculate *k_2_*, and subsequently, *k_1_* was calculated from the non-linear regression analysis-derived *k_2_*/*k_1_*. The results are also summarized in Table 3. As evident from this, the addition of the SKIK peptide tag led to values of *k_1_* and *k_2_* that were 8.65-fold and 22.5-fold higher, respectively.

Since the RBS is the same in the two examined mRNA sequences, it seems unlikely that the affinity between the ribosome and mRNA changes significantly. Therefore, the increase in *k*_1_ resulting from the addition of SKIK peptide tag suggests an augmentation in the pool of ribosomes capable of initiating translation. Similarly, the increase in *k_2_* implies an acceleration in the rate of translation from initiation to termination. These kinetic analyses indicate that the addition of the SKIK peptide tag could enhance the ribosomal turnover rate. Regarding studies on kinetic analysis of translation, previous analyses have utilized the incorporation of radiolabeled amino acids into synthesized proteins^42^, and more recently, elongation rates per codon have been calculated based on ribosome density observed on mRNA through ribosome profiling ^43,44^. To our knowledge, our analysis, based on the production of sfGFP as an indicator, has not been reported previously and is a first attempt. Elongation rate per codon is commonly used as an indicator of translation efficiency, with reported values ranging from 1.66 codons/s^45^ and in *E. coli* in vivo to 10 codons/s^42^ in vitro, 4.2 codons/s for *S. cerevisiae,* and 5.2 codons/s for mouse stem cells^43^

In this study, SKIK-SecM AP-sfGFP gene used consists of 267 codons. The apparent elongation rate per codon calculated from the estimated *V_max_* of SKIK-SecM AP-sfGFP was 1.2 codons/s. Although this value is lower than those reported in previous studies, considering that ribosomal stalling by SecM is not completely alleviated by SKIK peptide tag ^39^, and our reaction model disregards the translation initiation and termination stages which require time, the calculated elongation rate appears reasonable. Although it should be noted that use of this model might be limited, we concluded that the simple translation rate analysis method proposed in this study worked well for our purpose.

### Effect of EF-P and SKIK peptide

EF-P is a translation factor reported to promote the synthesis of poly-Pro motif. Therefore, we investigated how the SKIK peptide tag, capable of relieving ribosomal stalling induced by poly-Pro, and EF-P, would affect the translation of sequences containing poly-Pro. We examined using four sequences including -5 aa, +0 aa, GGGG, and AAAA. Among these, the ribosomal stalling was most effectively alleviated by SKIK peptide tag alone in -5 aa (Figure 2A). As a result of EF-P addition at a final concentration of 1 µM, the protein production was increased to 1.51-fold and 4.40-fold in -5 aa and +0 aa, respectively, compared with no additive. In particular, the protein production was highest for -5 aa with EF-P, indicating a synergistic effect by SKIK peptide tag and EF-P (Figure 4A). It suggests the possibly that EF-P and SKIK peptide tag individually promote translation in this case. By contrast, GGGG and AAAA, which do not contain SKIK, did not show any change in protein levels with addition of EF-P. To confirm the nascent state of SKIK peptide is essential for translation promotion, CFPS of four mRNA templates was conducted with the addition of the chemically synthesized free SKIK peptides at 100 µM which was 100 times higher concentration of ribosome in CFPS reaction. As expected, no influence was observed (Figure 4B). Huter et al. demonstrated that EF-P recognizes the P-site tRNA and the E-site mRNA codon, stabilizing the conformation of the P-site tRNA and thereby promoting peptidyl bond formation^46^. Taken together with the results of additional of free SKIK peptide and our previous study^39^, it is likely that nascent state of SKIK peptide tag is crucial for translation promotion. Therefore, we expect that the mechanism of action of externally functioning EF-P and SKIK peptide tag at the nascent state may be different. Since the mechanism of translation enhancement by SKIK peptide tag has not yet been elucidated, we believe that further structural analysis will be necessary.

**Figure 4.**
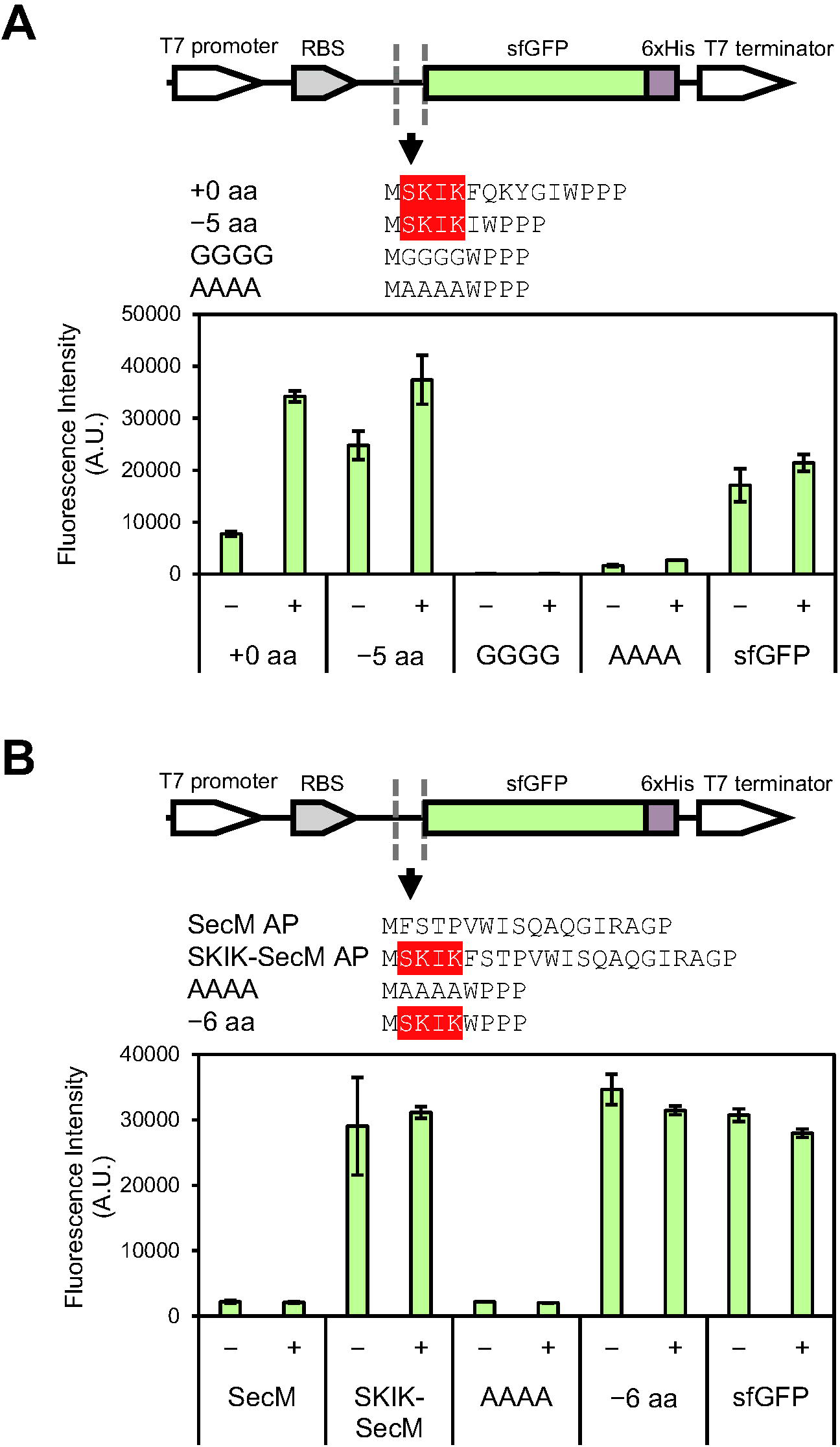
Effect of EF-P and free SKIK peptide on protein production in CFPS. A) Fluorescence intensity of sfGFP by in vitro translation using mRNA (1.61 µM) after 90 min reaction. +/− indicates the presence/absence of EF-P (1 µM). B) Fluorescence intensity of sfGFP by in vitro translation using mRNA after 90 min reaction. The mRNA concentration was 1.19 µM except for SecM AP and SKIK-SecM AP (2.37 µM). +/− indicates the presence/absence of synthesized free SKIK peptide (100 µM).

## Conclusions

In this study, we revealed that the translation enhancing SKIK peptide tag effectively alleviates ribosomal stalling induced by the WPPP when the distance between them is closer in both the PURE cell-free protein synthesis and in vivo protein production systems, resulting in significant increment of production of proteins containing such poly-Pro motif. A kinetic analysis based on the simple translation model showed that the ribosome turnover is enhanced by more than 10-fold when the SKIK peptide tag. Furthermore, we found that the use of both SKIK peptide tag and EF-P synergistically enhanced translation for proteins containing poly-Pro. Our findings and simple strategy have practical significance in enhancing protein production and contribute to the advancement of synthetic biology.

## Methods

### DNA primer list

DNA primers used in this research are summarized in Supplementaly Table 1.

### Plasmid construction

Total 21 plasmids were prepared by whole plasmid PCR using our previously constructed pET22b-SKIK-WPPP-sfGFP-His as the template and DNA primers listed in Supplementary Table 1. *Dpn*I treated PCR products were purified with silica column (FastGene Gel/PCR Extraction Kit, Nippon Genetics Co., Ltd., Tokyo, Japan) and then HiFi assembly reaction (New England Biolabs, Ipswich, MA) was carried out to connect each end of linear DNA products. The plasmids were prepared using plasmid mini prep kit (FastGene Plasmid Mini Kit, Nippon Genetics Co., Ltd.) from overnight cultured *E. coli* DH5alpha transformants. The sequences of all plasmids confirmed by Sangar sequencing are available in Supporting Information.

### Cell-free protein synthesis (CFPS)

DNA fragments containing T7 promoter and terminator used for CFPS were prepared by PCR with the primer pair F1 and R1, constructed plasmids as the template, and Tks Gflex DNA polymerase (98°C 1 min; (98°C 10 sec; 50°C 5 sec; 68°C 30 sec) × 30 cycles; 68°C 1min; 12°C ∞, Takara, Kusatsu, Japan). Amplified DNA was purified using silica column and used for in vitro mRNA synthesis. For in vitro mRNA synthesis, the T7 RiboMAX Large-Scale RNA Production System (Promega, Madison, WI, USA) was used and the synthesized mRNAs were purified with NucleoSpin RNA Plus (Takara). The concentration of mRNA was measured by Nano Drop One (Thermo Fisher scientific, MA, USA) and the molecular weight were calculated using RNA Molecular Weight Calculator on AAT Bioquest (https://www.aatbio.com).

CFPS reactions were carried out with PURE*frex* 2.1 (GeneFrontier Corporation, Kashiwa, Japan) under the following condition. Solution I, 4 µL; Solution II, 0.5 µL; Solution III (20 μM ribosome), 0.5 µL; 10 mM cysteine, 0.5 µL; RNase Inhibitor (TOYOBO, Osaka, Japan), 1 µL; at final reaction volume of 10 µL; incubated at 37°C for 90 min. The final concentration of the template mRNA depends on the examination. An elongation factor EF-P (GeneFrontier Corporation) and synthesized SKIK peptide dissolved in nuclease-free water (94.6% purity, GenScript, Tokyo) was added to achieve a final concentration of 1 µM and 100 µM, respectively, as needed.

### In vivo expression

Terrific broth (Bacto tryptone, 12 g; yeast extract, 24 g; KH_2_PO_4_, 6.8 g; Na_2_PO_4_・12H_2_O, 7.1 g; MgSO_4_, 0.15 g; (NH_4_)_2_SO_4_, 2.58 g; 10%(w/v) glucose, 5 mL; and 8%(w/v) lactose; 25 mL, per 1 L, adjusted pH 7.4) was used as the autoinduction medium. The single colonies of ECOS *E. coli* BL21(DE3) (Nippon Gene, Tokyo, Japan) transformed with each plasmid were inoculated to 4 mL of LB liquid and grown with shake at 37°C for 19 h. Then the aliquots (50 µL) were transferred to the 1 mL of terrific broth and cultured at 30°C until the OD600 reached 2.3–6.9. All media were supplemented with 100 µg/mL ampicillin. After collecting the bacterial cells by centrifugation and washing with phosphate buffer saline buffer (PBS, 137 mM NaCl, 2.7 mM KCl, 10 mM Na_2_HPO_4_, 1.8 mM KH_2_PO_4_; pH 7.4), the OD600 values of each sample were adjusted to the same. Protein was extracted from the cells using *E.coli*/Yeast Protein Extraction Buffer (Tokyo Chemical Industry Co., Ltd., Tokyo, Japan), and the lysate fraction was used for analysis.

### Preparation of Standard sfGFP

ECOS *E. coli* BL21(DE3) was transformed with pET22b-sfGFP-His and cultured in 100 mL of LB medium at 37°C with shake until the OD600 value reached 0.4∼0.6. IPTG was then added at a final concentration of 0.1 mM and cultured at 37°C for another 5 h. Bacterial cells were collected by centrifugation (4°C, 9000 rpm, 10 min), resuspended in 10 mL of PBS, and then the bacteria were crushed by sonication (30 min). The supernatant of the lysate was applied to a Ni-NTA agarose HP column (Fujifilm Wako Pure Chemical Corporation, Osaka, Japan) equilibrated with Binding buffer (20 mM sodium phosphate buffer, 0.5 M NaCl, 20 mM imidazole, pH 8.0), and the column was washed 3 times with 6 mL of binding buffer and eluted 6 times with 1 mL of Elution buffer (20 mM sodium phosphate buffer, 0.5 M NaCl, 500 mM imidazole, pH 8.0). The eluate was added to the dialysis membrane and dialyzed for a total of 24 h (buffer was changed twice during the dialysis). The concentration of purified sfGFP was calculated from the A280 nm value (A.U.) measured by Nano Drop One. Molecular weight (27788.3 Da) and molar extinction coefficient (19,035 M^-1^cm^-1^) of sfGFP were calculated using ProtParam tool on Expasy (https://www.expasy.org)^47^.

### Fluorescence measurement

Each CFPS reaction solution was diluted 25-fold with water and dispensed 50 µL per well into Flat Bottom Microfluor Plates Black (Thermo Fisher scientific, MA, USA), and the fluorescence intensity of sfGFP was measured by a microplate reader (Infinite 200 PRO, TECAN, ZH, Switzerland) at excitation wavelength 485 nm (bandwidth 9 nm)/emission wavelength of 535 nm (bandwidth 20 nm).

### Real-time monitoring of the translation in CFPS

The reaction was performed using a real-time PCR system (StepOnePlus Real-Time PCR System, Life Technologies, CA, USA) by incubating at 37°C, and the fluorescence intensity was measured every minute after 30 s. Fluorescence intensity per sfGFP molecule was calculated using purified sfGFP (14.6 µM, 1.46 µM, 0.146 µM, 0.0146 µM, and 0.00146 µM) as the standard. Translation rate defined as the generated sfGFP per ribosome per time was calculated from the rate of the fluorescence increase where the slope was the greatest and linear for all samples.

### Western Blotting

The samples (1 µL) were subjected to SDS-PAGE and transferred to a nitro cellulose membrane (Bio-Rad Laboratories, CA, USA). The membrane was blocked using 3% skim milk in PBST (137 mM NaCl, 2.7 mM KCl, 10 mM Na_2_HPO_4_, 1.8 mM KH_2_PO_4_, 0.05 % Tween-20; pH 7.4) for 1 h at room temperature. After washing three times with PBST, the membrane was incubated with Anti-His-tag mAb-HRP-DirectT (MBL, Tokyo, Japan) at 1:5000 dilution with Can Get Signal Solution 2 (Toyobo, Osaka, Japan) for 1 h at room temperature. After washing with PBST three times, blots were visualized with TMB solution (Nacalai tesque, Kyoto, Japan).

## Supporting Information

The Supporting Information including DNA sequences for plasmid with GenBank format and additional experimental data (PDF) is available free of charge.

## Supporting information

sequences of plasmid DNA

## Author contributions

Y.N., H.N., T.K., and T.O.K. designed the study and analyzed the data. Y.N. performed CFPS experiments. R.F. performed in vivo expression experiments. Y.N. and T.O.K. wrote the manuscript. T.O.K. supervised the project.

## Acknowledgment

This research was supported by Japan Science and Technology agency FOREST Program (grant No. JPMJFR2204), KAKENHI 23K04989, and GteXProgram Japan Grant Number JPMJGX23B6 and JPMJGX23B4. PURE*frex* 2.1 and supplemental components used in this study were provided by GeneFrontier Corporation. We thank Mrs. Kana Yamauchi (Nagoya University) for her help to the preparation of the purified sfGFP. We also thank Prof. Hisashi Hemmi (Nagoya University) and Prof. Tomokazu Ito (Nagoya University) for their kind advice and help on translation analysis.

## Abbreviations

RAP: Ribosomal arrest peptide
RBS: ribosome binding site
PTC: peptidyl-transferase center
CFPS: cell-free protein synthesis
sfGFP: superfolder green fluorescent protein

## Notes

### Competing Interest Statement

The authors have declared no competing interest.

### Summary of Updates

Author information updated; English revised; Supplemental Information uploaded; Minor corrections of Figures provided

