## Supplemental Information for "Effect of translation enhancing nascent SKIK peptide on the arrest peptides containing consecutive Proline"

Supplementary Table 1. DNA primers used in this study.

| Name | sequence (5′→3′) |
| --- | --- |
| G1_F_v2 | AAATAAAAGGTTTCCAGAAGTACGGGATTTGGCC |
| G1_R_v2 | TCTGGAAACCTTTTATTTTAGACATATGTATATCTCCTTCTTAAAG |
| G2_F | AAATAAAAGGTGGATTCCAGAAGTACGGGATTTGGCC |
| G2_R | CTTCTGGAATCCACCTTTTATTTTAGACATATGTATATCTCCTTCTTAAAG |
| G3_F | GGTGGAGGTTTCCAGAAGTACGGGATTTGGCC |
| G3_R_v2 | CTTCTGGAAACCTCCACCTTTTATTTTAGACATATGTATATCTCCTTCTTAAAG |
| G4_F | ATAAAAGGTGGAGGTGGCTTCCAGAAGTACGGGATTTGG |
| G4_R | GCCACCTCCACCTTTTATTTTAGACATATGTATATCTCC |
| G5_F | GGTGGAGGTGGCGGATTCCAGAAGTACGGGATTTGGCC |
| G5_R | GAATCCGCCACCTCCACCTTTTATTTTAGACATATGTATATCTCCTTCTTAAAG |
| G6_F | GGTGGAGGTGGCGGAGGTTTCCAGAAGTACGGGATTTGGCC |
| G7_F | GGTGGAGGTGGCGGAGGTGGCTTCCAGAAGTACGGGATTTGGCC |
| G7<_R | ACCTCCGCCACCTCCACCTTTTATTTTAGACATATGTATATCTCCTTCTTAAAG |
| G8_F | GGTGGAGGTGGCGGAGGTTTCCAGAAGTACGGGATTTGGCC |
| G9_F | GGTGGAGGTGGCGGAGGTGGCTTCCAGAAGTACGGGATTTGGCC |
| G10_F | GGTGGAGGTGGCGGAGGTGGCGGTGGAGGATTCCAGAAGTACGGGATTTGGCC |
| G11_F | GGTGGAGGTGGCGGAGGTGGCTTCCAGAAGTACGGGATTTGGCC |
| G12_F | GGTGGAGGTGGCGGAGGTGGCGGTGGAGGAGGTGGTTTCCAGAAGTACGGGATTTGGCC |
| G13_F | GGTGGAGGTGGCGGAGGTGGCGGTGGAGGAGGTGGTGGATTCCAGAAGTACGGGATTTGGCC |
| F_Fw | AAAATAAAACAGAAGTACGGGATTTGGCCGC |
| FQ_Fw | AAAATAAAAAAGTACGGGATTTGGCCGCCC |
| FQK_Fw | AAAATAAAATACGGGATTTGGCCGCCCCC |
| FQKY_Fw | AAAATAAAAGGGATTTGGCCGCCCCCTG |
| FQKYG_Fw | AAAATAAAAATTTGGCCGCCCCCTGCAAG |
| FQKYGI_Fw | AAAATAAAATGGCCGCCCCCTGCAAGTAAA |
| F_Rv | GTACTTCTGTTTTATTTTAGACATATGTATATCTCCTTCTTAAAG |
| FQ_Rv | CCCGTACTTTTTTATTTTAGACATATGTATATCTCCTTCTTAAAG |
| FQK_Rv | AATCCCGTATTTTATTTTAGACATATGTATATCTCCTTCTTAAAG |
| FQKY_Rv | CCAAATCCCTTTTATTTTAGACATATGTATATCTCCTTCTTAAAG |
| FQKYG_Rv | CGGCCAAATTTTTATTTTAGACATATGTATATCTCCTTCTTAAAG |
| FQKYGI_Rv | GGGCGGCCATTTTATTTTAGACATATGTATATCTCCTTCTTAAAG |
| G_replce_ | ATGGGTGGCGGAGGGTGGCCGCCCCCTGCAAGTAAAG |
| G_replce_Rv | CCACCCTCCGCCACCCATATGTATATCTCCTTCTTAAAGTTAAACAAAATTATTTC |
| A_replace_F | ATGGCAGCTGCCGCGTGGCCGCCCCCTGCAAGTAAAG |
| A_replace_R | CCACGCGGCAGCTGCCATATGTATATCTCCTTCTTAAAGTTAAACAAAATTATTTC |
| F1 | ATCTCGATCCCGCGAAATTAATACG |
| R1 | TCCGGATATAGTTCCTCCTTTCAG |

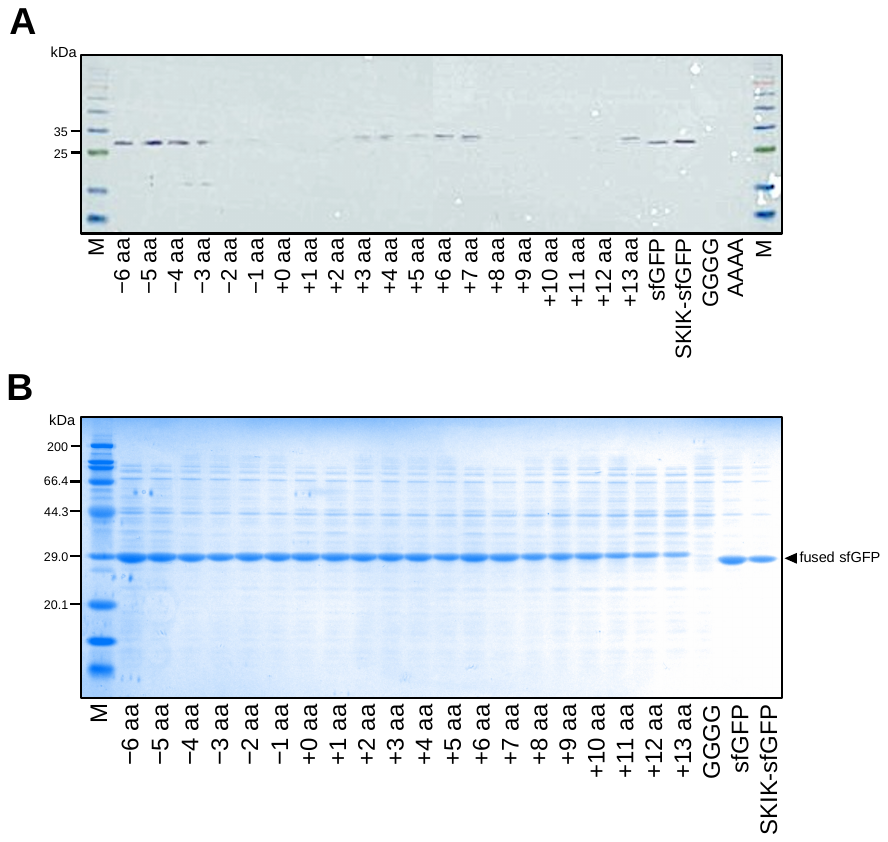

**Supplementary Figure 1. SDS-PAGE analysis of in vitro and in vivo protein production.**

A) The full image of Western Blotting in Figure 2A. The markers (M) used were ExcelBand 3-color Regular Range Protein Marker (SMOBiO, Taiwan, Hsinchu).
B) The full image of CBB staining result in Figure 2C. M indicates protein marker (Protein Molecular Weight Marker (Broad), Takara).

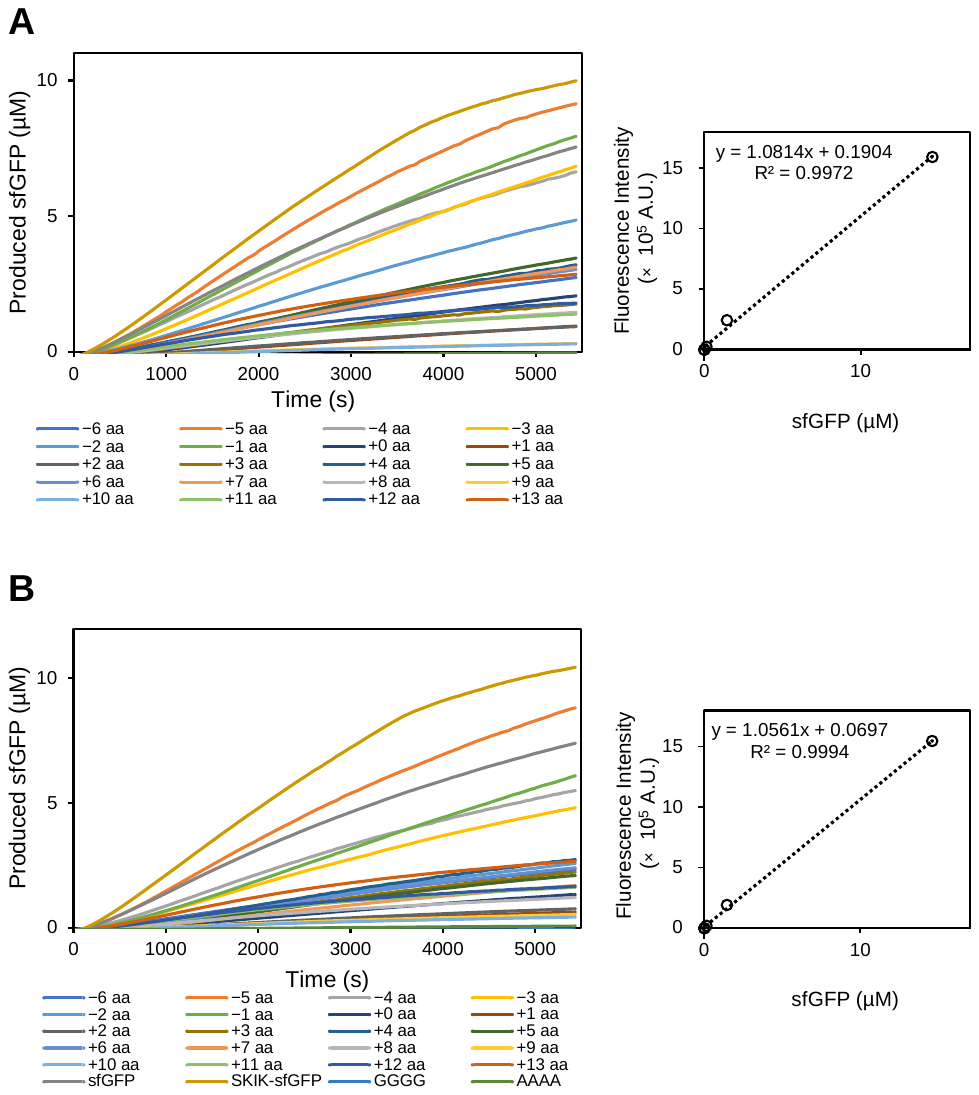

**Supplementary Figure 2. Real-time monitoring of the produced sfGFP in an in vitro translation.**

The sample names and sequences are corresponding to those of Figure 1. Fluorescence intensity of the produced sfGFP from mRNA was monitored during 90min (5,400 s) CFPS reaction to evaluate the influence of the distance between the SKIK peptide tag and WPPP poly-Pro motif. Two independent monitoring results (left) and standard curves of the purified sfGFP (right) are shown as A and B. Translation rate in Figure 2B was calculated based on the rate of increase in sfGFP in the 450–990 s.
